# Genes responsible for avoiding attack of a beetle, relating to the duration of death feigning

**DOI:** 10.1101/2021.05.13.443969

**Authors:** Keisuke Tanaka, Ken Sasaki, Kentarou Matsumura, Shunsuke Yajima, Takahisa Miyatake

## Abstract

Predator avoidance is an important behavior that affects the degree of adaptation of organisms. We compared the DNA variation of one of the predator-avoidance behaviors, the recently extensively studied “death-feigning behavior,” between the long strain bred for feigning death for a long time and the short strain bred for feigning death for a short time. To clarify how the difference in DNA sequences between the long and short strains corresponds to the physiological characteristics of the death-feigning duration at the transcriptome level, we performed comprehensive and comparative analyses of gene variants in *Tribolium castaneum* strains using DNA-re-sequence. The duration of death feigning involves many gene pathways, including caffeine metabolism, tyrosine metabolism, tryptophan metabolism, metabolism of xenobiotics by cytochrome P450, longevity regulating pathways, and circadian rhythm. Artificial selection based on the duration of death feigning results in the preservation of variants of genes in these pathways in the long strain. When an animal wake up from a near-death experience is closely related to its success in avoiding predation. This study suggests that many metabolic pathways and related genes may be involved in the decision-making process of anti-predator animal behavior by forming a network in addition to the tyrosine metabolic system, including dopamine, revealed in previous studies.

## Introduction

Since Edmund^1^, much research has focused on the behaviors adopted by animals to avoid attack by enemies^2^. Death feigning (or thanatosis, tonic immobility, playing possum, playing dead, post-contact immobility, and so on) that have recently received special attention is one way to avoid enemy attack^3,4,5,6^. It has also been considered an adaptive behavior for females to avoid male harassment^7,8,9,10,11^ and for individuals to avoid worker aggressions in social insects^12^. Although the adaptive significance of death-feigning behavior has become widely recognized, very little research has been done on its molecular mechanisms.

Recently, Uchiyama et al.^13^ compared transcriptomes of beetle strains selected for short and long durations of death feigning. In *Tribolium castaneum*, strains divergently selected for short (S strains) and long (L strains) durations of death feigning, which is activated by external stimuli, have been established in the laboratory^3,14,15,16^.

A previous study identified 518 differentially expressed genes (DEGs) between the strains by transcriptome analysis, because RNA sequencing (RNA-seq), is rapidly gaining momentum in an effort to reveal the molecular mechanisms underlying physiological mechanisms^13^. The study revealed that tyrosine metabolic pathways including dopamine synthesis genes, stress-response genes, and insulin signaling pathways were differentially activated between individuals of short and long strains^13^. However, we cannot determine which part of the DNA caused the degree of transcription to change by transcriptome analysis alone.

A reciprocal crossing experiment between short and long strains for duration of death feigning showed that it occurred more frequently and for shorter periods in the F1 population with dominance in the short direction. From the F2 population, the death-feigning duration showed continuous segregation, indicating the duration of death feigning is controlled by polygenes in *T. castaneum*^17^. This result is consistent with the finding of many expressed RNAs involved by Uchiyama et al.^13^. The duration of death feigning has been found to be multilaterally expressed with other traits of insects: for example, locomotor activity in *T. castaneum*^14^, *T. confusum*^18^, and *T. freemani*^19^, flight ability in *Callosobruchus chinensis*^20^, life history traits in *C. chinensis*^21^, and mating behavior in *T. castaneum*^22^ and *C. chinensis*^23^. Therefore, it is easy to predict that the duration of a behavioral trait will be genetically affected by many other traits. Thus, we need to clarify how many genes influence the selection for the duration of death feigning on a DNA level.

To clarify how differences in DNA sequences between the long and short strains correspond to the physiological characteristics of the death-feigning duration at the transcriptome level, we performed comprehensive and comparative analyses of gene variants in *T. castaneum* strains using DNA-resequencing.

## Results

The results of resequencing analysis showed variations of DNA sequence from the reference sequence in both long and short strains, and the variations were detected more frequently in the long strain in a whole genome. The variations were present in several genes including caffeine metabolism, tyrosine metabolism, tryptophan metabolism, metabolism of xenobiotics by cytochrome P450, longevity regulating pathway, and circadian rhythm.

Small nucleotide variants as SNV and multi-nucleotide variants (MNV) as deletion, insertion, and replacement were detected in long and short strains. The numbers of small variants in total were larger in long strains than short strains (Fig. 1). The most frequent type of small variants was SNV, and the proportions of SNV were 86.3% (5,813 / 6,734) in long strains and 86.7% (1,279 / 1,476) in short strains, respectively (Fig. 1A). The SNVs compared with the reference nucleotide occurred frequently between adenine and guanine or cytosine and thymine in both long and short strains (Fig. 1B), and the frequencies were up to three times as large as other base combinations, indicating more frequent transition and fewer transversion variants. Deletion and insertion ranged from one to nine bases in both long and short strains, with one and three bases especially were frequently deleted or inserted (Fig. 1C). Homozygosity presented more frequently than heterozygosity in all linkage groups, and it was approximately two-fifteenths times and two-twelfths times as large as heterozygosity in long and short strains, respectively (Fig. 1D). Homozygosity of variants was the most frequent in linkage group 7 (LG7) in the long strain and in linkage groups 2 (LG2) and 8 (LG8) in the short strain, respectively. The ratios of homozygosity to heterozygosity were the largest in LGX and LG2 in long and short strains, respectively.

**Figure 1.**
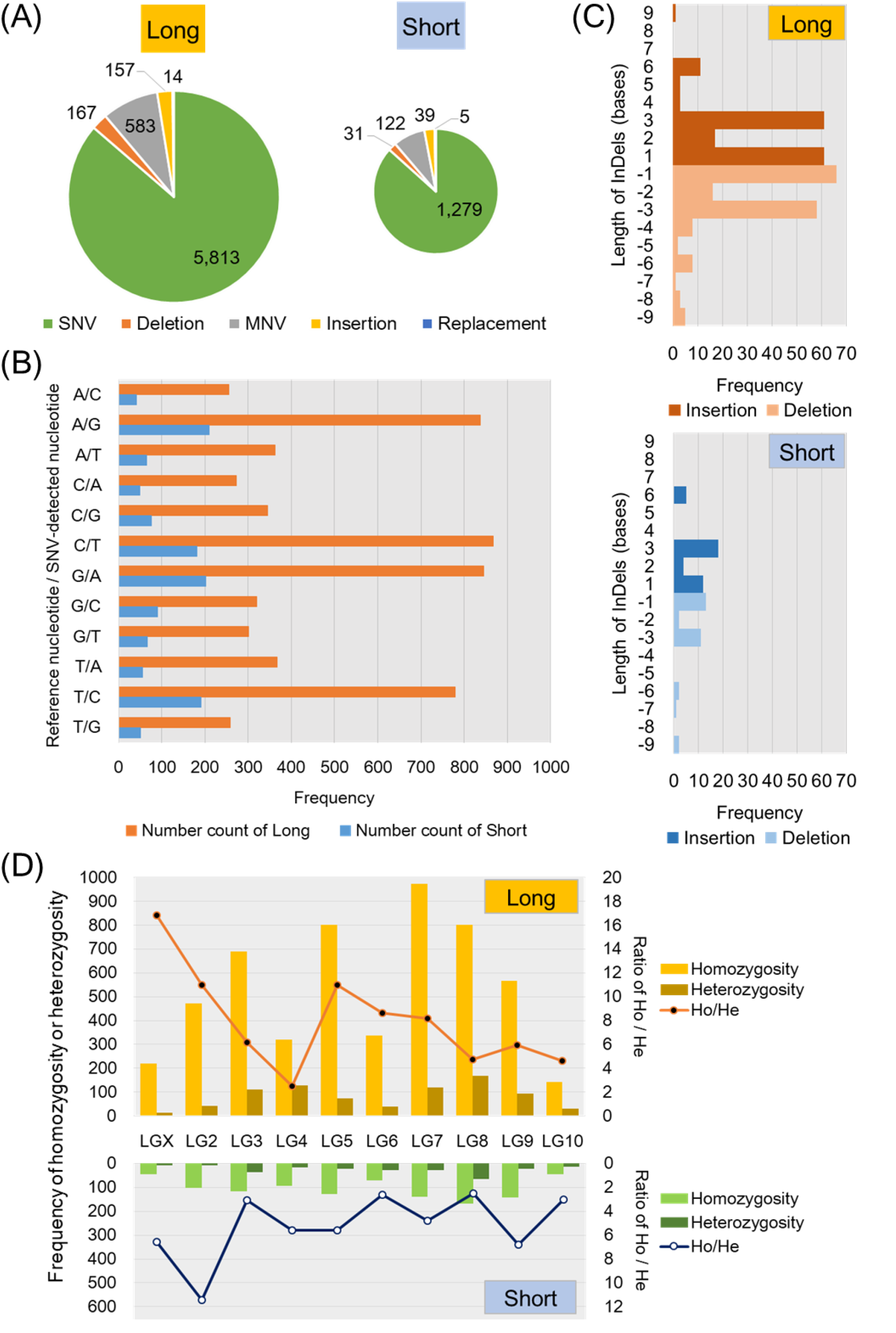
Analytical results of small variants of DNA sequence in long and short strains. Proportion of small variants as SNV, MNV, deletion, insertion, and replacement in long and short strains (A). The numbers of small variants are indicated as the diameter of a pie graph. Frequencies of the SNVs in both long and short strains were compared with the reference nucleotide (B). Insertion and deletion ranged from one to nine bases in both long and short strains (C). Frequency of homozygosity or heterozygosity in all linkage groups in long and short strains (D).

Genes with variants were more numerous in the long strain (3,384) than the short strain (1,075), and 718 genes were overlapped between the strains (Fig. 2A). Among these genes, the most frequent number of non-synonymous variants per gene was 1 in both strains, and the frequency gradually decreased as the number increased (Fig. 2B). The functions of genes with variants were sorted into four categories by enrichment analyses (Fig. 2c). In the biological process, molecular function, and Kyoto Encyclopedia of Genes and Genomes (KEGG) pathway, the long strain had a larger fold enrichment and larger statistical values than the short strain, while the short strain had a larger fold enrichment and statistical values in the “cellular component” than the long strain. The Gene Ontology (GO) term of “metabolic process” had the largest statistical value in the long strain, and KEGG Ontology (KO) terms “ECM-receptor interaction”, “ABC transporters” and “other glycan degradation” had larger statistical values in the long strain than the short strain.

**Figure 2.**
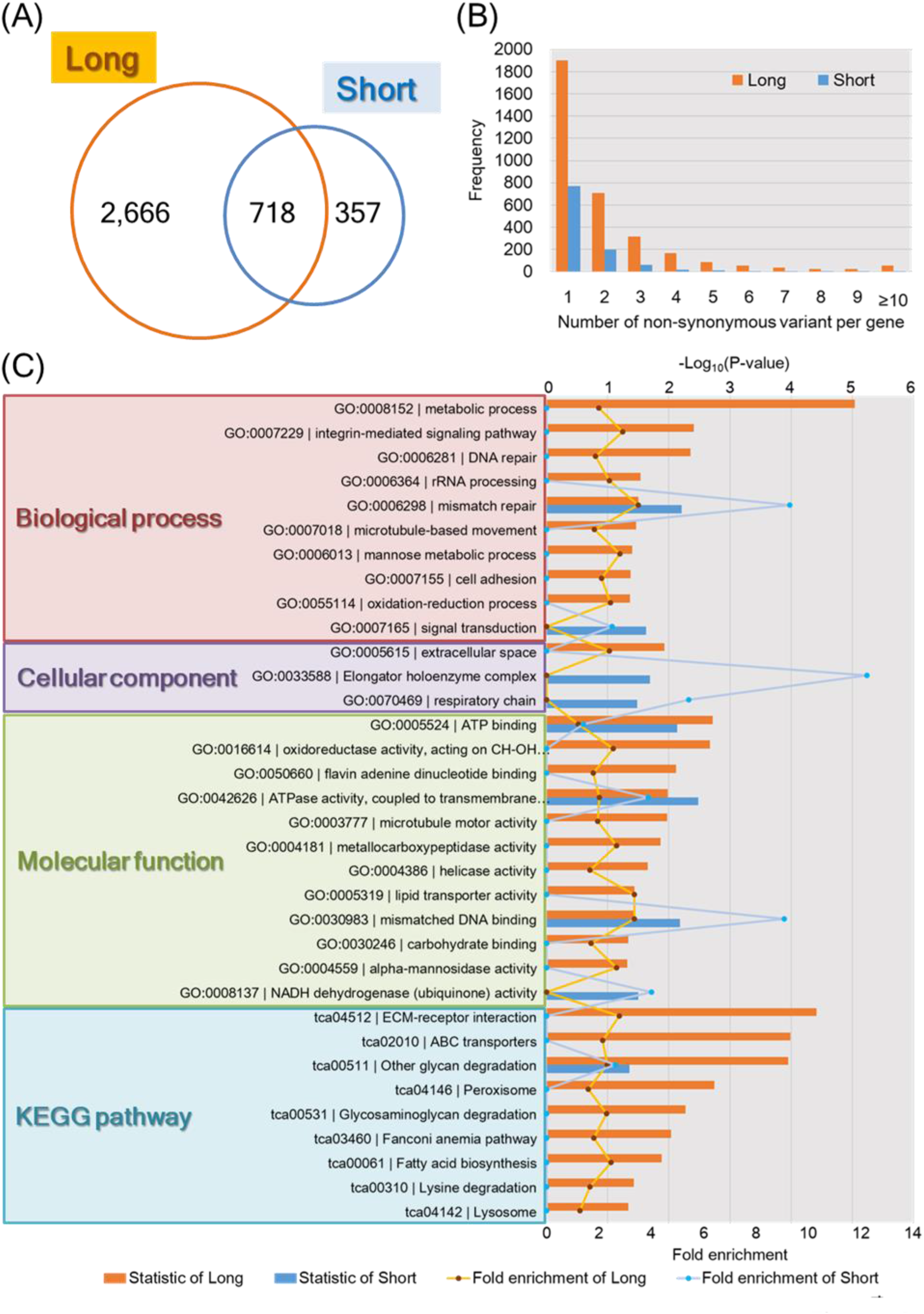
Analytical results of small variants in genes. Numbers of genes with variants in long and short strains (A). Frequency of the number of non-synonymous variants per gene in long and short strains (B). Enrichment analyses of the function of genes with variants sorted into four categories (biological process, cellular component, molecular function, and KEGG pathway) (C).

Structural variations including large-scale InDel, copy number variation (CNV), and presence/absence variation (PAV) were analyzed in both strains (Fig. 3). Large-scale insertions and deletions were analyzed in 10 to 390 bases, and 15–20 bases of insertion and deletion were the most frequent in both strains (Fig. 3A). CNV deletions were present more frequently in sizes ranging from 5 to 16 kbases than in other size scales that we examined in both strains, whereas CNV duplications were constantly less frequent at 0 to 3 cases (Fig. 3B). In a larger size scale of nucleotides, up to 7000 kbases, the presence of variations less than 500 kbases of nucleotide sizes was most frequent in the long strain (Fig. 3C). All of these are illustrated on each linkage group in Fig. 4A, indicating large-scale insertions and deletions constantly appearing in each linkage group (A and B), less frequent CNV duplications (C), and more frequent CNV deletions (D). Large CNV deletions were present in LG6 and LG7 in the long strain and in LG2 in the short strain, respectively. These variations were sorted into GO and KO terms (Fig. 4B). The term of “neuroactive ligand - receptor interaction” had the largest statistical value in the long strain.

**Figure 3.**
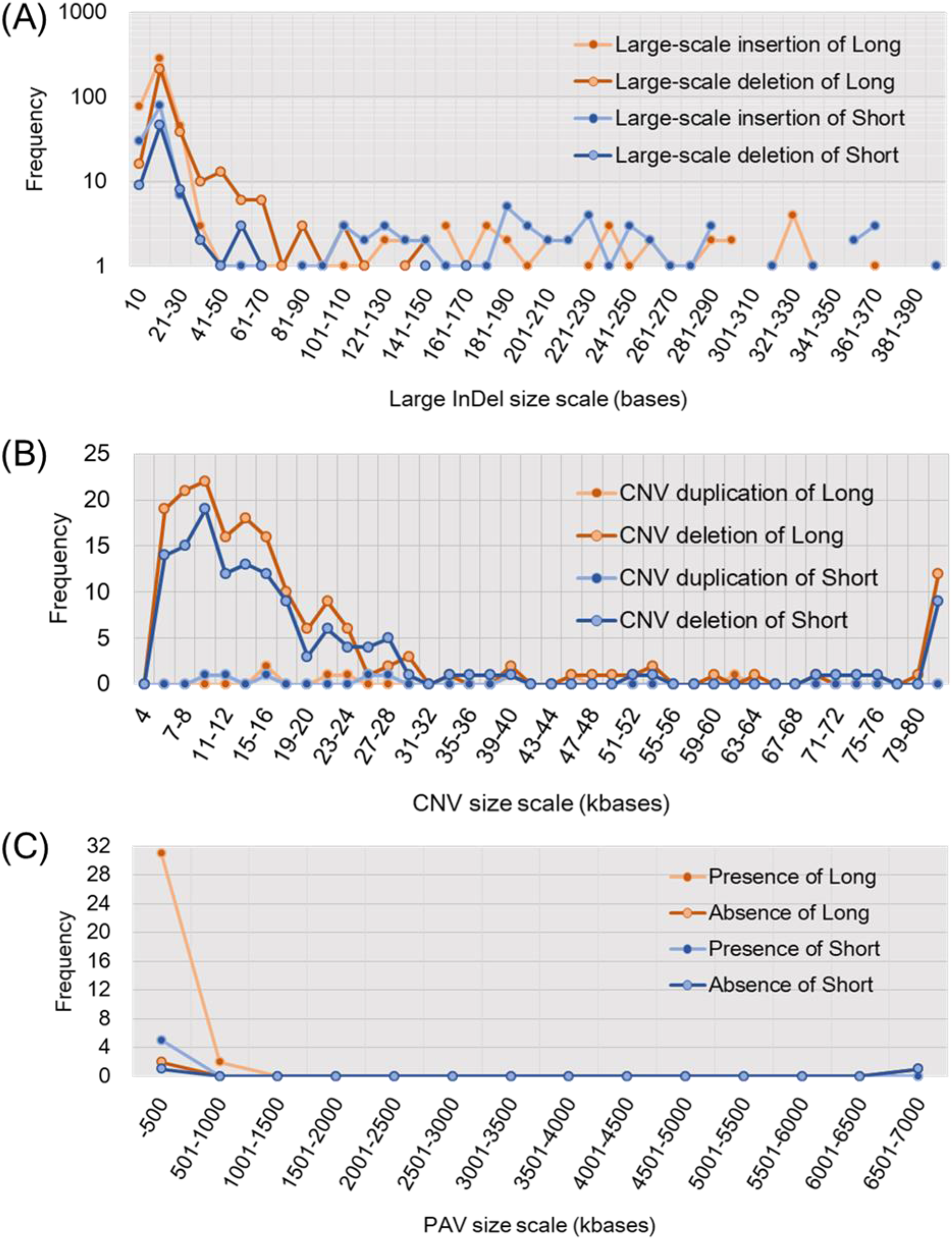
Structural variations including large-scale insertion and deletion (InDel) (A), copy number variation (CNV) (B), and presence/absence variation (PAV) (C) in long and short strains. Large-scale InDels were analyzed from 10 to 390 bases. CNV duplication and deletions were analyzed were from 4 to 80 kbases. PAV were analyzed from 0 to 7000 kbases.

**Figure 4.**
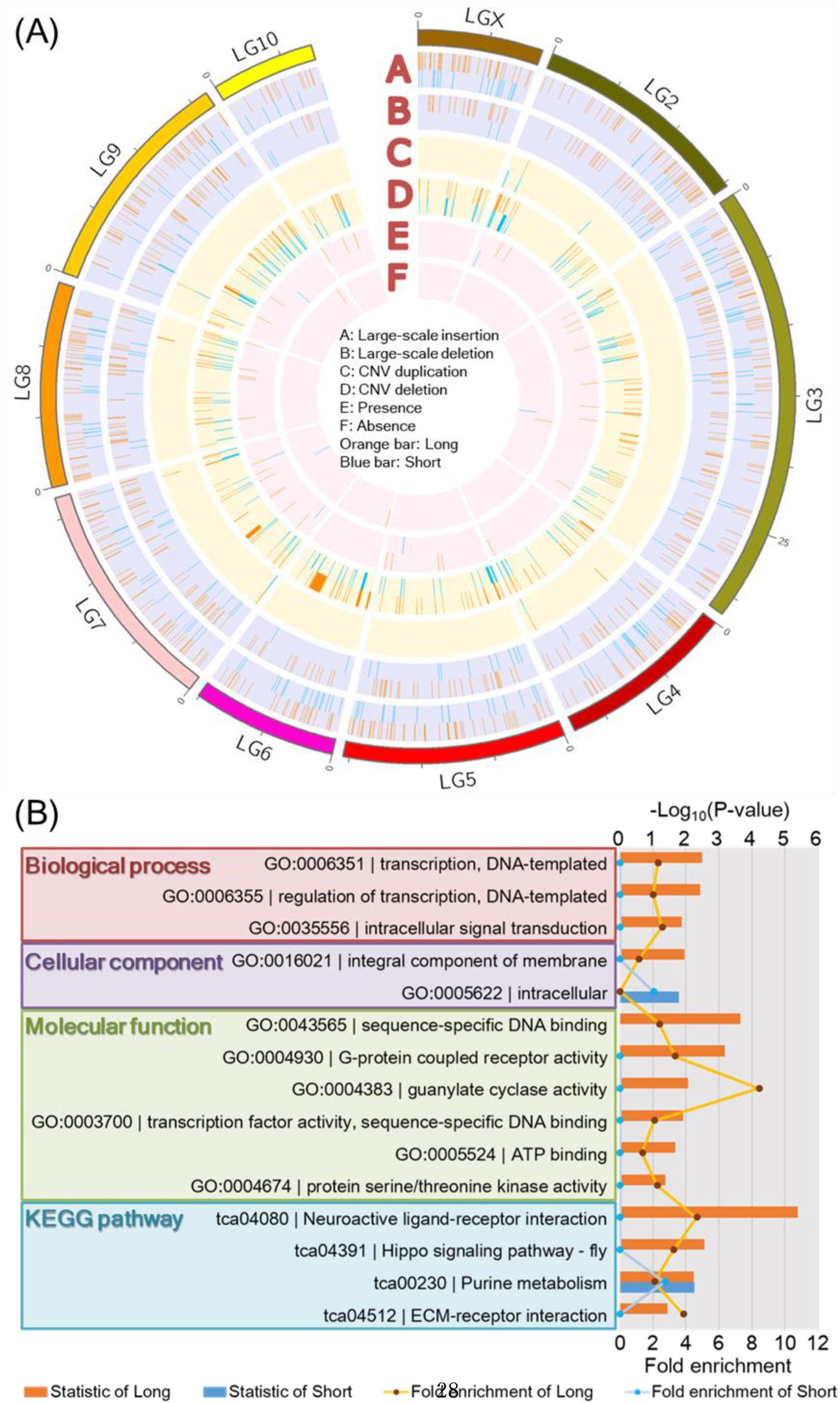
Structural variations on each linkage group in long and short strains (A). Structural variations include large-scale insertion and deletion, CNV duplication and deletion, and presence and absence of variation. The long and short strains are indicated by orange and blue lines, respectively. GO and KO terms are from function of genes with variants (B).

A protein – protein interaction (PPI) network including enzymes involved in dopamine metabolism was constructed (Fig. 5). Tyrosine hydroxylase (*Th*) was connected with DOPA decarboxylase (*Ddc*) and dopamine *N*-acetyltransferase (*Dat*), and these enzymes have been reported as differentially expressed genes in the long strain analyzed by RNA-seq^13^. *Th* also had variations of DNA sequence in the short strain (Fig. 5). Among the PPI network, proteins with variants were more frequent in the long strain. Yellow-like protein had variants in the long strain, and it was indirectly connected with *Ddc* and *Th* and directly with *Dat*.

**Figure 5.**
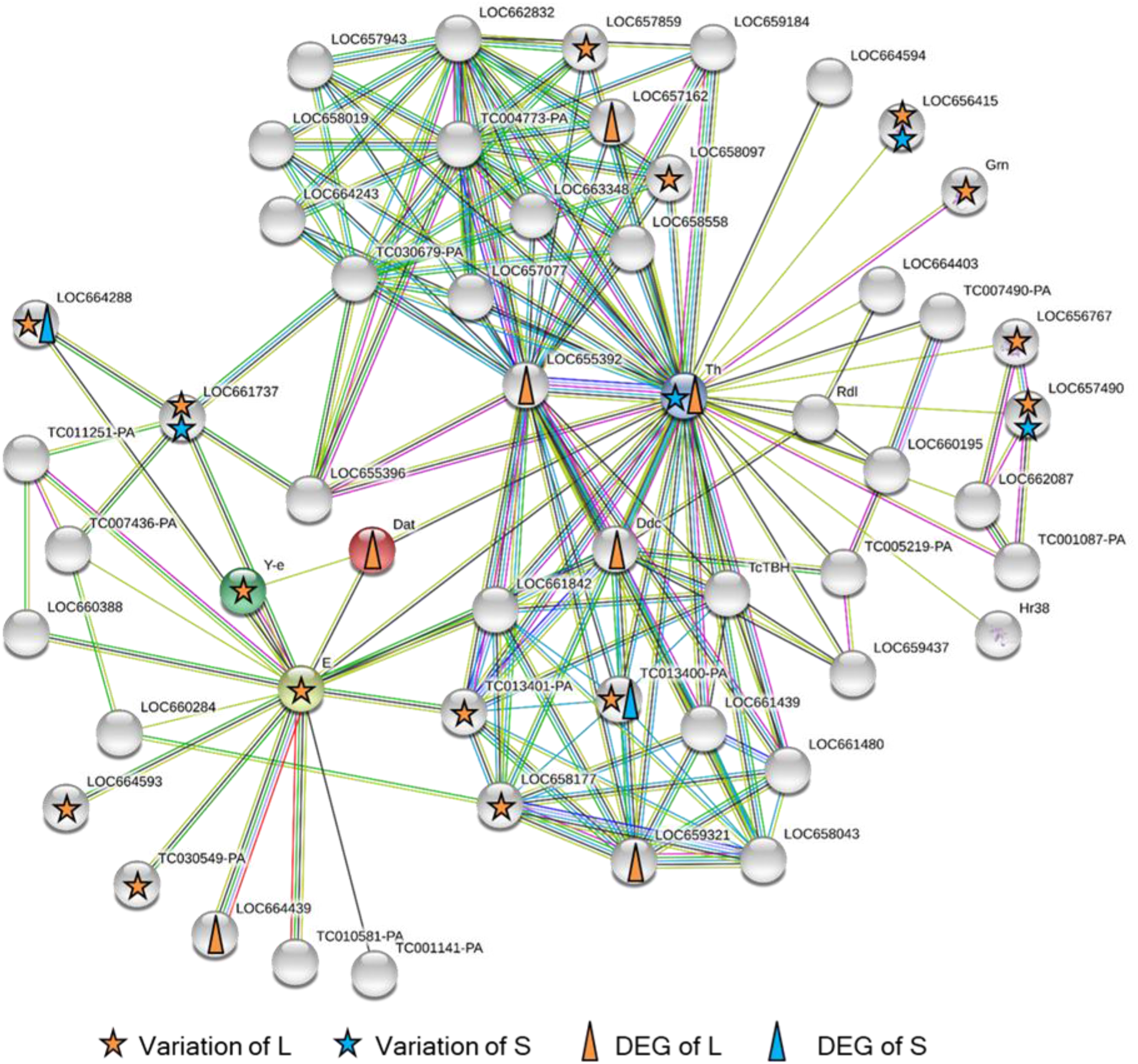
Protein-protein interaction (PPI) network including enzymes involved in dopamine metabolism. Lines indicate the relationships between genes. Stars and triangles indicate genes with variants and differentially expressed genes (DEGs), respectively.

Pathways containing genes with variants in both long and short strains were analyzed in “caffeine metabolism (tca00232)” (Fig. 6A), “tyrosine metabolism (tca00350)” (Fig. 6B), “tryptophan metabolism (tca00380)” (Fig. 6C), “metabolism of xenobiotics by cytochrome P450 (tca00980)” (Fig. 6D), “longevity regulating pathway - multiple species (tca04213)” (Fig. 6E), and “circadian rhythm - fly (tca04711)” (Fig. 6F). Tyrosine metabolism and longevity-regulating pathways have been listed as pathways containing focal genes with different expressions between long and short strains as detected by RNA-seq^13^. Except for circadian rhythm, the numbers of variants of genes in these pathways were larger in the long strain than the short strain (Fig. 6).

**Figure 6.**
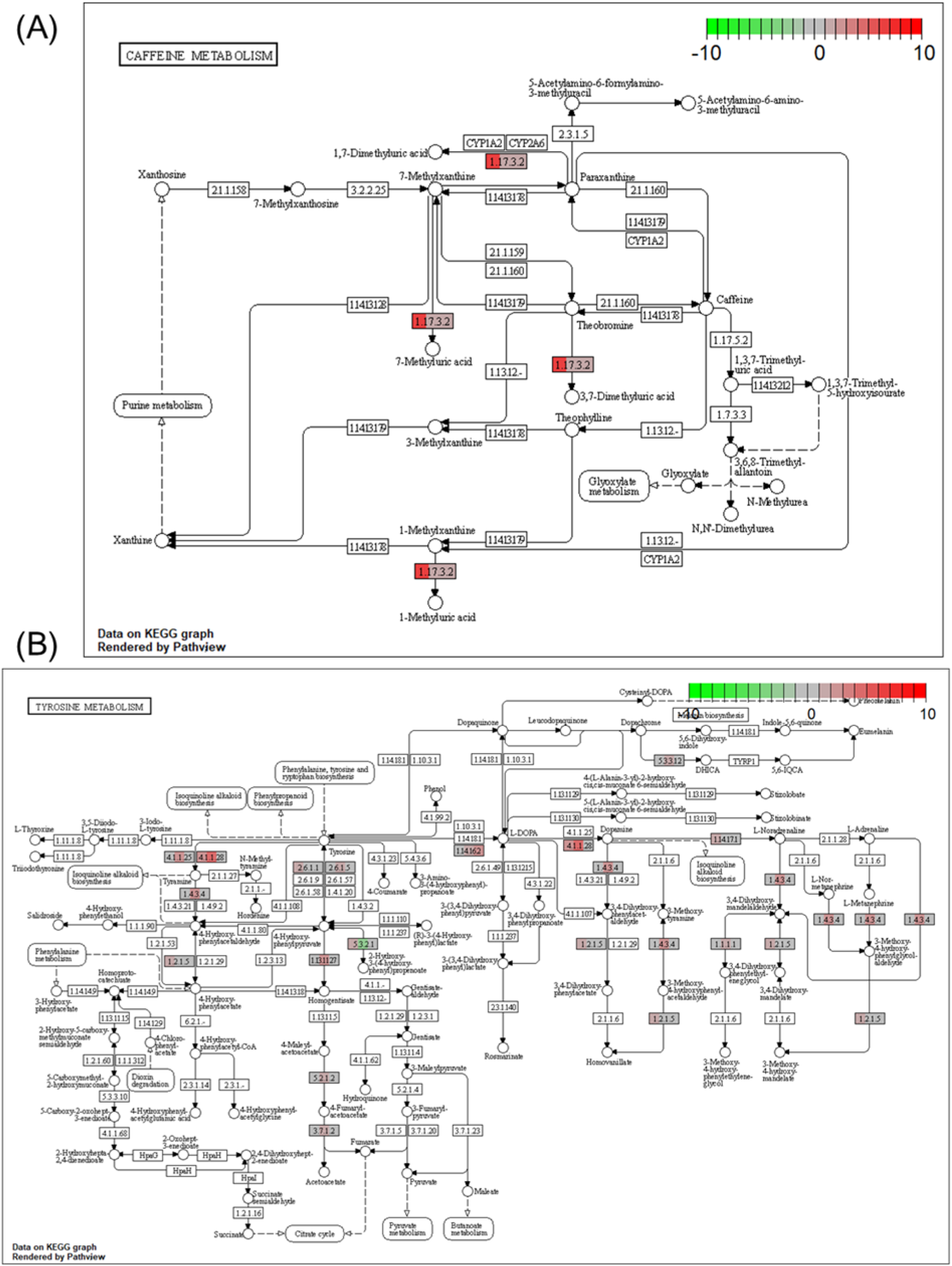
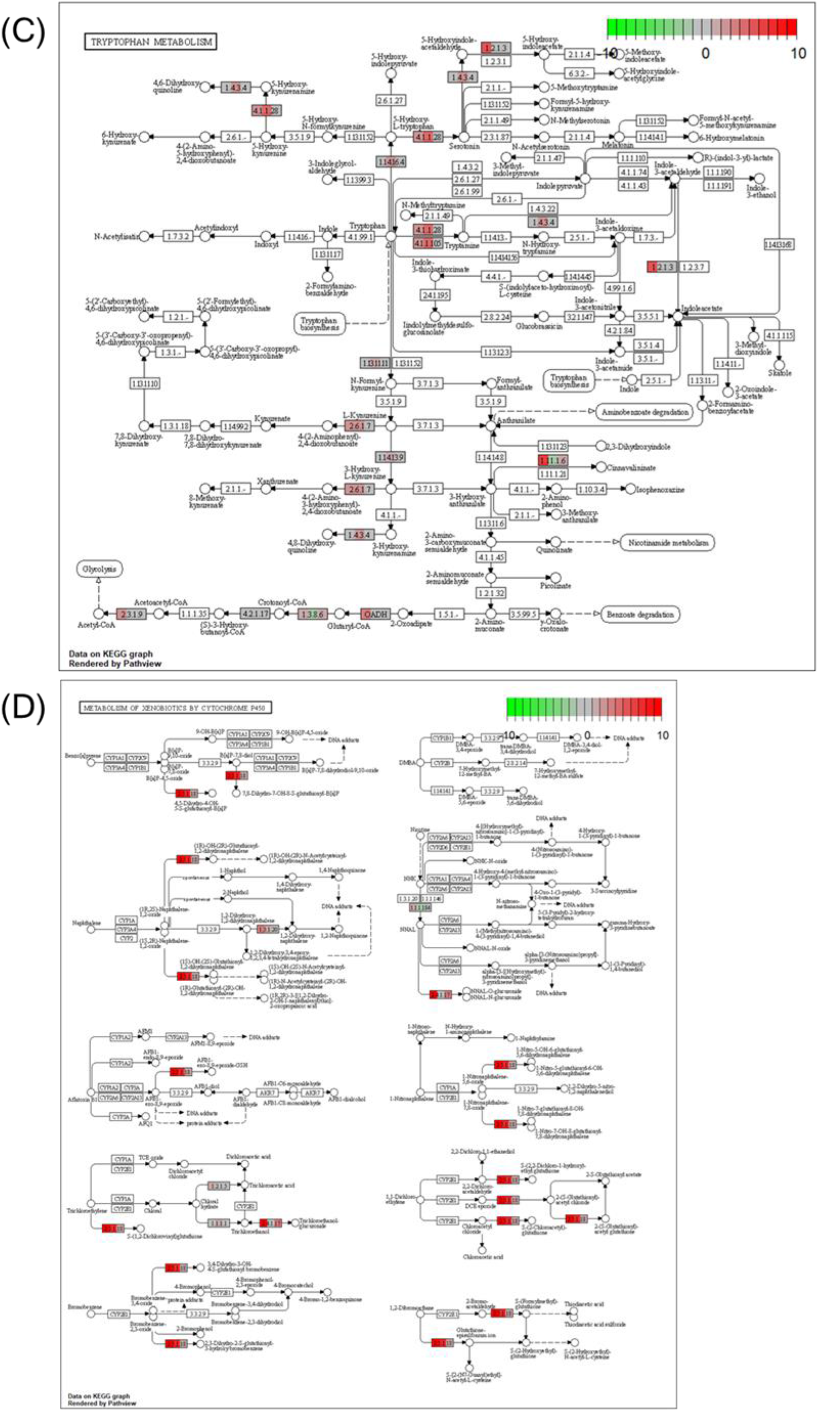
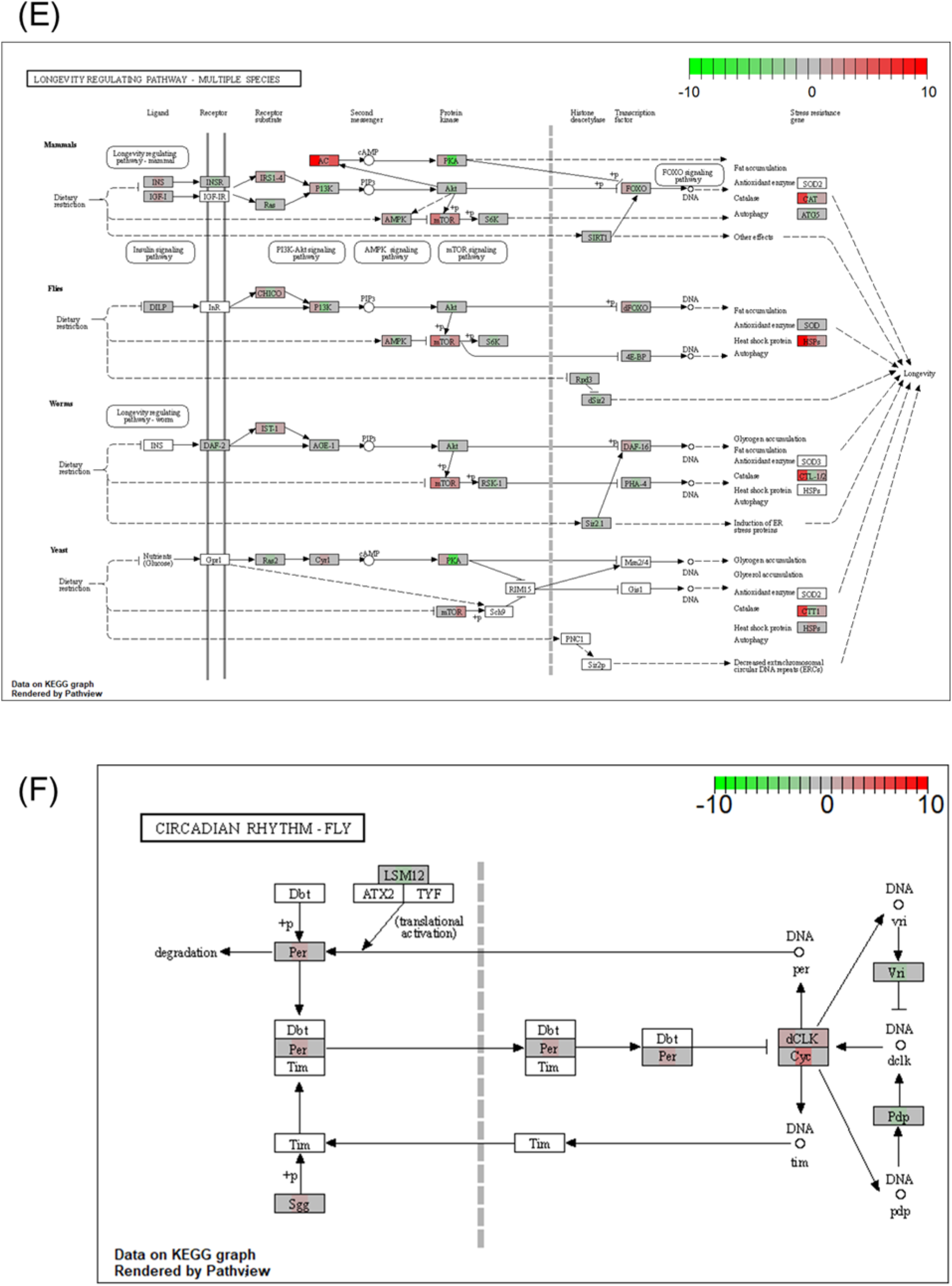
Functional genes with frequency of variants in KEGG pathways. Caffeine metabolism (A), tyrosine metabolism (B), tryptophan metabolism (C), metabolism of xenobiotics by cytochrome P450 (D), longevity regulating pathway (E) and circadian rhythm (F) are indicated.

## Discussion

The present study compared DNA sequences in a whole genome between the long strain and reference individuals or the short strain and reference individuals in *T. castaneum*. The analyses detected small variants, including SNV, MNV, deletion, insertion and replacement in particular genes in the particular metabolic and signaling pathways in both long and short strains. Especially, the long strain had more variants in a particular gene than those in the short strain. This was the case in the structural variations including large-scale InDel, CNV, and PAV. These large-scale variations were frequently detected in the long strain. The frequent variations of DNA sequences in the long strain may cause production of more diverse gene expressions in the long strain. Artificial selection based on the duration of death feigning results in the preservation of variants in genes in these particular metabolic and signaling pathways in the long strain, whereas the short strain with fewer variants of the DNA sequence might be selected more strongly than the long strain.

### Caffeine metabolism

In the KEGG pathway, there were significant inter-strain differences in CYOIA2, 7-methyluric acids, 3,7-dimethyluric acid, and 1-methyluric acid (Fig. 6A). Oral ingestion and injection of caffeine have been shown to shorten the death-feigning duration of adults in the long strains of *T. castaneum*^24^. As similar phenomenon has been found in the related species *T. confusum*, in which strains selected for longer duration had shorter death-feigning duration when they had ingested caffeine orally^25^. Nakayama et al.^25^ concluded that the dopaminergic system plays an important part in controlling the genetic correlation between death feigning and activity levels in these *Tribolium* species. Caffeine is known as a dopamine activator. Therefore, the present result demonstrates the relationship between the caffeine-dopamine system and death feigning at the genome level.

### Tyrosine metabolism

The results of the transcriptome by the RNA-seq study^13^ and the current DNA analysis are consistent with the tyrosine metabolic system (Fig. 6B). Variants in DNA sequence were found in four genes in the long strain and one gene with variants in two sequence locations in the short strain. One of the four genes in the long strain was “*Ddc*”, which is involved in the direct synthesis of dopamine, and one of the genes in the short strain was “*Th*”, which is involved in the synthesis of L-dopa, the precursor of dopamine. In the previous RNA-seq study^13^, the expression levels of these genes were found to be significantly higher in the long strain, so they may be affected by the DNA sequences revealed in the present results. The tyrosine and phenylalanine metabolic systems might be associated with the tryptophan metabolic system. Genome analysis showed more variants in genes of the tryptophan metabolic system than in the tyrosine metabolic system, but the proteins of the tryptophan metabolic system are related to the tyrosine metabolism. Because many variants seem to occur in the gene region related to glucose metabolism, the difference in the amount of ATP produced as a result of glucose metabolism may also affect the effects of caffeine. Insect hormone biosynthesis may also be related to this pathway. In the KEGG, there is a significant difference in franesol, juvenile hormone, and CYP307A1/2. The ecdysteroids may have an effect on tyrosine metabolism and dopamine synthesis. One of insect hormones is juvenile hormone synthesis, which may also be related to dopamine synthesis.

### Tryptophan metabolism

In the KEGG pathway of tryptophan metabolism, 16 genes showed differences between strains (Figure 6C). There are four possible explanations for how mutations in the tryptophan metabolic system indirectly affect gene expression. (1) Differences in serotonin levels and serotonin synthase gene expression may be related to the tryptophan metabolism. (2) The amount of melatonin, a metabolite of serotonin, and together with serotonin, is related to variations in circadian rhythms (also see Figure 6F), which will be discussed later. (3) Metabolism of the ommochrome system (ommochrome is a pigment substance that produces the color of the compound eye (individual eye)), and may be related to this metabolic system in relation to vision. (4) Synthesis of tryptophan into proteins that are needed (tryptophan is an amino acid, so it is a raw material for proteins). To prove these hypotheses, we need to conduct further studies at physiological and/or molecular levels.

### Metabolism of xenobiotics by cytochrome P450

Eight genes differed between strains (see Figure 6D). At phenotypic level, comparisons of metabolisms between the strains have not been conducted in *T. castaneum*. On the other hand, in *Callosobruchus chinensis*, long-lines selected for longer duration of death feigning exhibited higher rates of emergence, laid bigger eggs compared with strains selected for shorter duration of death feigning and greater reproductive effort, and also had a tendency to develop faster^22^. These changes in life-history traits, correlated with the duration of death feigning, may be related to metabolism in insects because strains with longer duration of death feigning can conserve energy to reproduce during their lives. This trade-off between anti-predator strategies and energy conservation might relate to the metabolism. Therefore, it is required to measure differences between the metabolisms of long and short strains of *T. castaneum*.

### Longevity regulating pathway

Differences in expression in the longevity regulating pathway were also found in an RNA-seq study^13^. This pathway contains a catalase gene (*CAT*) seen in a larger number of variants in the long strain. Kiyotake et al.^26^ has reported that the longer strains had lower relative expressions of *CAT*. This lower expression of *CAT* might be involved in the preservation of a large number of *CAT* variants in the long strain. Because *Tribolium* species have long longevity (more than 200 days), no comparison of lifetimes between strains was made. On the other hand, in *Callosobruchus chinensis*, long strains selected for longer duration of death feigning exhibited greater longevity compared short strains selected for shorter duration of death feigning and greater reproductive effort, and also had a tendency toward faster development^21^.

### Circadian rhythm

Long strain beetles have a significantly larger variation than short strain beetles in the three circadian genes including *Per* (period), *Sgg* (Shaggy), and *Cyc* (Cycle). Although the previous RNA-seq study showed the expression of *Dbt* (doubletime) is only slightly between the short and long strains (Uchiyama et al. 2019), the present DNA re-sequence did not find any difference in the *Dbt* gene between the strains. Therefore, it will be very interesting to compare the circadian rhythm of beetles derived from long and short strains in the future.

### Other genes

There are other differences in the DNA sequences of the long and short strains; for example, the glutathione metabolism pathway and stress-resistance and heat-shock genes. Using the same strains as the present study, Kiyotake et al.^26^ showed that the longer strains had lower relative expression of catalase (*CAT*) and growth-blocking peptide (*GBP*) genes compared to the short strains. These genes are related to anti-stress capacity, and Kiyotake et al.^26^ showed that the long-strain beetles are significantly more sensitive to environmental stressors such as mechanical vibration and high or low temperatures than the short-strain beetles. The difference in stress resistance between the strains may relate to stress-resistance genes and the heat-shock system.

## Conclusion

The duration of death feigning is related to many gene pathways, including caffeine metabolism, tyrosine metabolism, tryptophan metabolism, metabolism of xenobiotics by cytochrome P450, longevity regulating pathway, and circadian rhythm. Artificial selection based on the duration of death feigning results in the preservation of variants in genes in these pathways in the long strain. An animal’s decision on when to wake up from a near-death experience is closely related to its success in avoiding predation^5,27,28^. This study suggests that many metabolic pathways and related genes may be involved in the decision-making process of anti-predator animal behavior by forming a network in addition to the tyrosine metabolic system including dopamine revealed in previous studies.

## Materials and Methods

### (1) Insects

The red flour beetle, *Tribolium castaneum* (Herbst 1797), is a stored-product insect found worldwide and a model genome species, designated by the *Tribolium* Genome Sequencing Consortium^29^. The protocol for artificial selection for the duration of death feigning was described in Miyatake et al.^3^ Briefly, the duration of death feigning was measured in 100 male and 100 female adult beetles that were randomly selected. From these populations, 10 males and 10 females with the shortest and longest durations of death feigning were allowed to reproduce for the next generation. The selection regime was continued for more than 20 generations^14,15,16^.

### (2) DNA extraction

Female individuals in short or long strains were frozen by liquid nitrogen. Head and thoracic tissues without legs were removed from the frozen bodies by a pair of fine spring scissors. Each tissue was homogenized by the scissors and an electric homogenizer (T10+S10N-5G, IKA Works, Staufen, Germany) in an extraction buffer from an ISOGEN kit (Nippongene, Tokyo, Japan) according to the manufacturer’s instructions. The quality and quantity of the extracted DNA were determined at 230, 260, and 280 nm using a spectrophotometer (NanodropTM 2000, Thermo Fisher Scientific, MA, USA).

### (3) Library construction and sequencing

A total of 1 – 10 ng genomic DNA was fragmented by shearing to an average fragment size of 300 bp using an Adaptive Focused Acoustics sonicator (Covaris, Woburn, MA, USA). After purification, the paired-end DNA library was constructed using a KAPA Hyper Prep kit (KAPA Biosystems, Wilmington, MA, USA). The fragmented DNA was end-repaired, dA-tailed, and ligated with the paired-end adapter according to the manufacturer’s instructions. The adapter - ligated DNA was amplified by 14 cycles of high-fidelity polymerase chain reaction (PCR) amplification. Library quality and concentration were assessed using an Agilent Bioanalyzer 2100 (Agilent Technologies, Waldbronn, Germany) and an Agilent DNA 1000 kit. In addition, the library concentration was precisely determined using a KAPA Library Quantification Kit (Kapa Biosystems).

The paired-end libraries were sequenced by 200 cycles (2 × 100 bp) using the HiSeq 2500 (Illumina, San Diego, CA, USA). Reads were generated in FASTQ format using the conversion software bcl2fastq2 (Illumina, version 2.18). We submitted the read data to the Read Archive of DDBJ (accession number DRA011837).

### (4) Read mapping to reference whole genome

A series of data analyses was processed using CLC Genomics Workbench 12 (Qiagen, Hilden, Germany). After adapter trimming and quality filtering, the clean read data were mapped to the reference genome of *T. castaneum* (Tcas5.2) that was obtained from the NCBI genome database (https://www.ncbi.nlm/nih.gov/). The mapping parameters were as follows: mismatch cost = 2, insertion cost = 3, deletion cost = 3, length fraction = 0.9, and similarity fraction = 0.9. After local realignment of the mapped reads, duplicate PCR reads were discarded.

### (5) Small variant detection

Variant calling based on single nucleotide variation (SNV) and insertion and deletion (InDel) was performed using the CLC Genomics Workbench built-in tool “Fixed Ploidy Variant Detection”. The calling parameters were as follows: ploidy = 2, required variant probability = 90.0, minimum coverage = 10, minimum frequency = 20, minimum central quality = 40, and minimum neighborhood quality = 30. Furthermore, high-quality variants were selected using QUAL = 80. Genes with non-synonymous substitutions in the selected variants were enriched as Gene Ontology (GO) and Kyoto Encyclopedia of Genes and Genomes (KEGG) ontology (KO) terms using the web-based tool “DAVID 6.8” (https://david.ncifcrf.gov)^30^. The enriched terms were statistically analyzed using the modified Fisher’s exact test (*p* < 0.05) contained in the tool.

### (6) Structural variant detection

Variant calling based on copy number variation (CNV) was performed using CNVnator version 0.3.3^31^. The calling parameters were as follows: size ≥ 4000, normalized RD ≥ 2 or ≤ 0.5, and E-value by *t*-test statistics < 0.05. In comparison, variant calling based on presence/absence variation (PAV) was run using the CLC Genomics Workbench built-in tool “InDel and Structural Variants”. The genes contained in the identified regions were enriched as GO and KO terms using the web-based tool “DAVID 6.8” The enriched terms were statistically analyzed using the modified Fisher’s exact test (*p* < 0.05) contained in the tool.

### (7) Functional annotation between variant and differentially expressed genes

We investigated whether resequencing data was relevant to previous transcriptome analysis data^13^. A protein–protein interaction (PPI) network was constructed using the String version 11.0^32^. The target protein searched for DAT having a central role in this study. The parameters were customized as follows for the output data: minimum required interaction score, 0.600; max number of interactors to show no more than 50 interactors (both 1st and 2nd shells). Pathway analysis was performed for “caffeine metabolism (tca00232)”, “tyrosine metabolism (tca00350)”, “tryptophan metabolism (tca00380)”, “metabolism of xenobiotics by cytochrome P450 (tca00980)”, “longevity regulating pathway - multiple species (tca04213)”, and “circadian rhythm - fly (tca04711)” using the R package “Pathview”^33^.

### (8) Pathway analysis

KEGG metabolic maps (https://www.kegg.jp/kegg/kegg1.html) based on FCs between the L and S strains were highlighted using the R/Bioconductor package “Pathview” to compare expression levels during death feigning^34, 35, 36^.

## Author contributions statement

The study was conceived by K.T., K.S., K.M. S.Y., and T.M. The experiments were designed by K.T., K.S., S.Y., and T.M., and these were performed by K.T., K.S., and K.M. The data was analyzed by K.T., K.S., and TM. The manuscript was written by K.T., K.S., and T.M. All authors approved the final version prior to submission.

## Ethical approval and informed consent

N/A because of insects.

## Data Availability

The datasets generated during and/or analyzed during the current study are available from the corresponding author on reasonable request.

## Acknowledgements

This work was supported by the Japan Society for the Promotion of Science Grant-in-Aid for Scientific Research Grants 18H02510 (to T.M., K.S.), and the Cooperative Research Grant of the Genome Research for BioResource, NODAI Genome Research Center, Tokyo University of Agriculture (to T.M., K.S., S.Y.).

## References

1. Edmunds M. Defence in Aminals. Longman, London. ISBN-10: 0582441323 (1974).

2. Ruxton G, Allen WL, Sherratt TN, Speed M. Avoiding Attack: The Evolutionary Ecology of Crypsis, Warning Signals, and Mimicry. Second Edition, Oxford, UK: Oxford University Press (2018).

3. Miyatake, T., Katayama, K., Takeda, Y., Nakashima, A., Sugita, A., Mizumoto, M. Is death-feigning adaptive? Heritable variation in fitness difference of death-feigning behaviour. Proceedings of Royal Society B 271, 2293–2296 (2004).

4. Humphreys, R., Ruxton, G.D. A review of thanatosis (death-feigning) as an antipredator behaviour. Behavioral Ecology and Sociobiology 72, 22. https://doi.org/10.1007/s00265-017-2436-8 (2018).

5. Sendova-Franks, A.B., Worley, A., Franks, N. R. Post-contact immobility and half-lives that save lives. Proceedings of Royal Society B https://doi.org/10.1098/rspb.2020.0881

6. Sakai, M. Death-Feigning in Insects: Mechanism and Function of Tonic Immobility (Entomology Monographs). Springer 1st ed. (2021).

7. Dennis DS, Lavigne RJ. Ethology of *Efferia varipes* with comments on species coexistence (Diptera: Asilidae). Journal of Kansas Entomological Society 49, 48–62 (1976).

8. Lawrence SE. Sexual cannibalism in the praying mantid, *Mantis religiosa*: a field study. Animal Behaviour 43, 569–583 (1992).

9. Shreeve TG, Dennis RLH, Wakeham-Dawson A. Phylogenetic, habitat, and behavioural aspects of possum behaviour in European Lepidoptera. Journal of Research Lepidoptera 39, 80–85 (2006).

10. Bilde T, Tuni C, Elsayed R, Pekár S, Toft S. Death feigning in the face of sexual cannibalism. Biology Letter 2, 23–25 (2006).

11. Khelifa R. Faking death to avoid male coercion: extreme sexual conflict resolution in a dragonfly. Ecology 98, 1724–1726 (2017).

12. van Veen JW, Sommeijer MJ, Aguilar Monge I. Behavioural development and abdomen inflation of gynes and newly mated queens of *Melipona beecheii* (Apidae, Meliponinae). Insectes Sociaux 46, 361–365 (1999).

13. Uchiyama H, Sasaki K, Hinosawa S, Tanaka K, Matsumura K, Yajima S, Miyatake T. Transcriptomic comparison between beetle strains selected for short and long durations of death feigning. Scientific Reports 9, 1004: DOI 10.1038/s41598-019-50440-5 (2019).

14. Miyatake T, Tabuchi K, Sasaki K, Okada K, Katayama K, Moriya S. Pleiotropic anti-predator strategies, fleeing and feigning death, correlated with dopamine levels in *Tribolium castaneum*. Animal Behaviour 75, 113–121 (2008).

15. Miyatake T, Nakayama S, Nishi Y, Nakajima S. Tonically immobilized selfish prey can survive by sacrificing others. Proceedings of Royal Society B 276, 2763–2767 (2009).

16. Matsumura K, Miyatake T. Responses to relaxed and reverse selection in strains artificially selected for duration of death-feigning behavior in the red flour beetle, *Tribolium castaneum*. Journal of Ethology 36, 161–168 (2018).

17. Matsumura, K., Miyatake, T. Polygene control and trait dominance in death-feigning syndrome in the red flour beetle *Tribolium castaneum*. (under review).

18. Nakayama S, Nishi Y, Miyatake T. 2010. Genetic correlation between behavioural traits in relation to death-feigning behaviour. Population Ecology 52, 329–335

19. Konishi K, Matsumura K, Sakuno W, Miyatake T. Death feigning as an adaptive anti-predator behaviour: Further evidence for its evolution from artificial selection and natural populations. Journal of Evolutionary Biology 33, 1120–1128. (2019).

20. Ohno T, Miyatake T. Drop or fly? Negative genetic correlation between death-feigning intensity and flying ability as alternative anti-predator strategies. Proceedings of the Royal Society B 274, 555–560 (2007).

21. Nakayama S, Miyatake T. Positive genetic correlations between life-history traits and death-feigning behavior in adzuki bean beetle (*Callosobruchus chinensis*). Evolutionary Ecology 23, 711–722. (2009a).

22. Nakayama S, Miyatake T. Genetic trade-off between abilities to avoid attack and to mate: a cost of tonic immobility. Biology Letters 6, 18–20 (2009b).

23. Nakayama S, Miyatake T. A behavioral syndrome in the Adzuki bean beetle: Genetic correlation among death feigning, activity, and mating behavior. Ethology 116, 108–112 (2010).

24. Nishi Y, Sasaki K, Miyatake T. Biogenic amines, caffeine and tonic immobility in *Tribolium castaneum*. Journal of Insect Physiology 56, 622–628 (2010).

25. Nakayama S, Sasaki K, Matsumura K, Lewis Z, Miyatake T. Dopaminergic systems as the mechanism underlying personality in a beetle. Journal of Insect Physiology 58, 750–755 (2012).

26. Kiyotake H, Matsumoto H, Nakayama S, Sakai M, Miyatake T, Ryuda M, Hayakawa Y. Gain of long tonic immobility behavioral trait causes the red flour beetle to reduce anti-stress capacity. Journal of Insect Physiology 60, 92–97 (2014).

27. Ishihara R, Matsumura K, Jones JE, Yuhao J, Fujisawa R, Nagaya N, Miyatake T. Arousal from death feigning by vibrational stimuli: comparison of Tribolium species. Journal of Ethology 39, 107–113 (2021).

28. Franks NR, Worley A, Sendova-Franks AB. Hide-and-seek strategies and post-contact immobility. Biology Letters doi.org/10.1098/rsbl.2020.0892 (2021).

29. Tribolium Genome Sequencing Consortium. The genome of the model beetle and pest *Tribolium castaneum*. Nature 452, 949–955 (2008).

30. Huang DW, Sherman BT, Tan Q, Collins JR, Alvord WG, Roayaei J, Stephens R, Baseler MW, Lane HC, Lempicki RA. DAVID Gene Functional Classification Tool: A novel biological module-centric algorithm to functionally analyze large gene list. Genome Biol. 2007 Sep 4;8(9):R183. (2007).

31. Abyzov A, Urban AE, Snyder M, Gerstein M. CNVnator: An approach to discover, genotype, and characterize typical and atypical CNVs from family and population genome sequencing. Genome Research 21, 974–984 (2011).

32. Szklarczyk D, Gable AL, Lyon D, Junge A, Wyder S, Hietra-Cepas J, Simonovic M, Doncheva NT, Morris JH, Bork P, Jensen LJ, von Mering C. STRING v11: protein-protein association networks with increased coverage, supporting functional discovery in genome-wide experimental datasets. Nucleic Acids Research, 47 D607–D613 (2019).

33. Luo W, Brouwer C. Pathview: an R/Bioconductor package for pathway-based data integration and visualization. Bioinformatics, 29, 1830–1831 (2013).

34. Kanehisa, M. and Goto, S.; KEGG: Kyoto Encyclopedia of Genes and Genomes. Nucleic Acids Res. 28, 27–30 (2000).

35. Kanehisa Furumichi, M., Tanabe, M., Sato, Y., and Morishima, K.; KEGG: new perspectives on genomes, pathways, diseases and drugs. Nucleic Acids Res. 45, D353–D361 (2017).

36. Kanehisa, M., Sato, Y., Furumichi, M., Morishima, K., and Tanabe, M.; New approach for understanding genome variations in KEGG. Nucleic Acids Res. 47, D590–D595 (2019).

